# Population genomics of the widespread African savannah trees *Afzelia africana* and *Afzelia quanzensis* (Caesalpinioideae, Fabaceae) reveals no significant past fragmentation of their distribution ranges

**DOI:** 10.1101/730911

**Authors:** Armel S.L. Donkpegan, Rosalía Piñeiro, Myriam Heuertz, Jérôme Duminil, Kasso Daïnou, Jean-Louis Doucet, Olivier J. Hardy

**Author notes:** **Corresponding author:** Armel S.L. Donkpegan.

## Abstract

Few studies have addressed the evolutionary history of tree species from African savannahs at large geographic scales, particularly in the southern hemisphere (Zambezian region). *Afzelia* (Fabaceae: Caesalpinioideae) contains economically important timber species, including two species widely distributed in African savannahs: *A. africana* in the Sudanian region and *A. quanzensis* in the Zambezian region. To characterize the population genetic diversity and structure of these two species across their distribution ranges, we used nuclear microsatellites (simple sequence repeats, SSRs) and genotyping-by-sequencing (GBS) markers. Six SSR loci were genotyped in 241 *A. africana* and 113 *A. quanzensis* individuals, while 2,800 and 3,841 high-quality single nucleotide polymorphisms (SNPs) were identified in 30 *A. africana* and 12 *A. quanzensis* individuals, respectively. Both species appeared to be outcrossing (selfing rate ~ 0%). The spatial genetic structure was consistent with isolation-by-distance expectations based on both SSR and SNP data, suggesting that gene dispersal is spatially restricted in both species (*b*_*Ld (SSR)*_= −*0.005* and −*0.007* and *b*_*Ld (SNP)*_= −*0.008* and −0.006 for *A. africana* and *A. quanzensis*, respectively). Bayesian clustering of SSR genotypes failed to identify genetic structure within species. In contrast, SNP data resolved intraspecific genetic clusters in both species, illustrating the higher resolving power of GBS at shallow levels of divergence. However, the clusters identified by SNPs revealed low levels of differentiation and no clear geographical entities. These results suggest that, although gene flow has been restricted over short distances in both species, populations have remained connected throughout the large, continuous Savannah landscapes. The absence of clear phylogeographic discontinuities, also found in a few other African savannah trees, indicates that their distribution ranges have not been significantly fragmented during past climate changes, in contrast to patterns commonly found in African rainforest trees.

## 1. INTRODUCTION

Studies on the population genetic structure of African trees have largely focused on rainforest species (Hardy et al., 2013; Daïnou et al., 2014 & 2016; Duminil et al., 2015; Ikabanga et al., 2017; Demenou et al., 2018; Monthe et al., 2018). In contrast, the evolutionary history of trees from the drier Sudanian and Zambezian regions, situated respectively North and South of the Guineo-Congolian rainforest (APPENDIX 5), is still largely undocumented. In these phytogeographic regions, trees occur in savannah, woodlands, dry forests or gallery forests, thus, in vegetation types that cover a wide range of density in tree cover. Therefore, we can expect that the response to climate change and gene flow in these vegetation types differs from those occurring in the rainforests. The climatic changes of the Pleistocene have had a significant impact on the savannah vegetation; however, they did not necessarily lead to fragmentation as usually assumed for the African rainforests (Maley, 1996). During the dry and cold glacial periods, savannahs expanded in the tropical regions in detriment of rainforest, which survived in fragmented refugia. At extreme latitudes the savannah lost ground to the advances of steppes or desert (Lioubimtseva et al., 1998). Conversely, during the humid interglacial periods, savannahs have been replaced by rainforests in the tropics, but were able to expand northwards and southwards at extreme latitudes (Quézel, 1965; Lézine, 1989; Waller & Salzmann, 1999; Salzmann et al., 2002; Vincens et al., 2006; Watrin et al., 2009). In the absence of evidence of past fragmentation, we may expect that widespread savannah trees exhibit only weak or no genetic discontinuities within species.

To our knowledge, only five savannah tree species have been genetically investigated in Africa using population genetics approaches at large scales. Three of the species occur in the Sudanian savannah (Northern Hemisphere): the shea tree *Vitellaria paradoxa* (Allal et al., 2011; Logossa et al., 2011), the African Mahogany *Khaya senegalensis* (Sexton et al., 2015), and the Locust Bean *Parkia biglobosa* (Lompo et al., 2018). The other two species exhibit a Sudano-Zambezian distribution (Northern and Southern Hemispheres): the baobab, *Adansonia digitata* (Tsy et al., 2009; Kyndt et al., 2009) and the Arabic gum species *Acacia senegal* (Odee et al., 2012; Lyam et al., 2018). Within the Sudanian savannah, weak genetic structure was detected in *K. senegalensis* and *A. digitata*, while moderate differentiation was found in *A. senegal*. For *V. paradoxa* and *P. biglobosa* significant genetic structure was detected in the Sudanian savannah, although in both cases large genetically homogeneus clusters were spread in central west Africa (Logossa et al., 2011; Lompo et al., 2018). Within the Zambezian domain, significant population genetic structure was detected for *A. senegal*, but not for *A. digitata*. Regional studies of *Syzygium guineense* (Zigelski et al., 2019; restricted to part of its Zambezian distribution) and *V. paradoxa* in Ghana revealed very weak genetic structure (Lovett et. al, 2000).

*Afzelia* (Fabaceae) is a palaetropical genus represented by seven species in Africa, including two savannah and four rainforest species, as well as one putative species which is currently poorly characterized (Brummit et al., 2007). The genus also harbours four species in South-East Asia (Donkpegan et al., 2014). The two African savannah species are widely distributed in Sub-Saharan Africa and occur in allopatry (Donkpegan et al., 2014): *Afzelia africana* Sm. ex Pers occurs in the Sudanian region (from Senegal to Sudan; Aubréville 1968; Geerling 1982) and *Afzelia quanzensis* Welw. in the Zambezian region (from southern Somalia to northern South Africa). Recently, it has been shown that the two savannah species are diploid, as opposed to the rainforest species, which are tetraploid (Donkpegan et al., 2015). In a recent phylogenetic study of African species of *Afzelia*, the genus was estimated to have emerged in open habitats (woodland and savannah) during the early to mid-Miocene (c. 20 to 14.5 Ma), whereas *A. quanzensis* and *A. africana* originated during the mid or late Miocene (c. 14.5 Ma to 8 Ma, Donkpegan et al., 2017). African *Afzelia* species are intensively logged for their timber (Donkpegan et al., 2014). However, the population genetic structure and evolutionary processes within the savannah species have not been investigated at a large geographic scales, despite the fact that genetic information may be useful for the development of sustainable management strategies for conservation and timber production (Lowe and Allendorf, 2010). Nuclear Simple Sequence Repeats (nSSR, also called microsatellites) markers revealed low genetic diversity in populations of *A. quanzensis* at small spatial scales, suggesting a limited evolutionary potential in this species (Jinga et al., 2016; Jinga & Ashley, 2018). In SE Asia large-scale population genetic studies have been performed on *A. xylocarpa* using SSR markers (Pakkad et al., 2009; Pakkad et al., 2014). In addition to genetic diversity, the spatial genetic structure between individuals or populations can inform on the evolutionary processes operating in a species and can thus be of interest for conservation management (Frankham et al., 2017). Among different metrics used to estimate relatedness between individuals (Frankham et al., 2017), the kinship coefficient is the most commonly used in tests for isolation by distance and to estimate the spatial extent of gene flow.

Population genetics studies in tropical trees have mostly used SSRs. Recent technological advances in high-throughput sequencing allow sequencing large portions of the genome in non-model organisms at a reasonable cost, thus offering increased resolution for the characterization of population genetic patterns and the inference of evolutionary processes (Ekblom & Galindo 2011). In this study, we use nuclear microsatellites (SSRs, Donkpegan et al., 2015) and single nucleotide polymorphisms (SNPs) derived from Genotyping by Sequencing (GBS) to investigate the population genomic processes in the two savannah species of *Afzelia* across their distribution ranges. This study addresses the following questions: (1) Does the genetic variation at large geographic scales reveal discrete gene pools and/or a pattern of isolation-by-distance within each species? (2) Do species show contrasting levels of genetic diversity and effective population size, or signatures of demographic change compatible with past bottlenecks and/or population growth? Our main objectives are to: (i) estimate the genetic diversity and population genetic structure of *A. africana* and *A. quanzensis*, using nuclear SSRs and SNPs, (ii) characterize the relatedness pattern between individuals in each species to test for isolation by distance, and (iii) understand the origin of these patterns using methods for demographic inference. Using SNP data on widespread savannah species, this paper is one of the first population genomic studies of tropical woodland trees distributed across western and southern Africa.

## 2. MATERIALS AND METHODS

### 2.1. Study species

*Afzelia africana* (Detarioideae, Fabaceae) occurs in the Sudanian region both in dry savannah and in dry forests (Aubréville, 1959, Ahouangonou et al., 1995; Gerard & Louppe, 2011). It can also occur in semi-deciduous forests, but at very low densities (Satabié, 1994). It has a wide ecological amplitude but it prefers areas with > 900 mm annual rainfall and grows at elevations of up to 1400 m and can reach up to 20 m in height. The fruiting period lasts six to eight months and fruits may persist on trees for the following six months (Bationo et al., 2001; Ouédraogo-Koné et al., 2008). *Afzelia quanzensis* occurs in the savannahs of Zambezian region, from Somalia to Angola and the north of South Africa. It has been reported in semi-deciduous coastal forests in Kenya (Brummitt et al., 2007) but also in dry forests, usually in deep sandy soils and also on rocky ridges (Jacana, 1997). The species is drought resistant but frost sensitive. It is a deciduous, medium to large-sized tree, 12-15 m high (reaching 35 m under ideal conditions, Coates-Palgrave, 2002). *Afzelia* species are hermaphrodite and pollinated by insects (e.g. bees, Kato et al. 2008; Ariwaodo and Harry-Asobara 2015). They have large dehiscent woody pods containing characteristic black and red seeds (Jacana, 1997; Gerhardt and Todd, 2009). Squirrels predate the seeds while monkeys, rodents (*Proechimys spp.*), and birds (mainly hornbills) act as dispersers (Van Wyk & Van Wyk, 1997; Gathua, 2000; Bationo et al., 2001; Gerard & Louppe, 2011).

### 2.2. Sampling and DNA extraction

Plant tissue samples were collected directly in the field or in herbaria (National Herbarium of the Netherlands (herbarium code WAG of the Index Herbariorum), the Botanical Garden of Meise (BR) and Université Libre de Bruxelles (BRLU) in Belgium), recording the geographic coordinates of individual sampling locations. Our sampling is representative of the known distribution ranges of the two species, in the Sudanian and Congolian biogeographic regions for *A. africana* and in the Somalian, Zambezian and South African regions for A. *quanzensis* (APPENDIX 5). We sampled 241 *A. africana* individuals from 41 West and Central African locations and 113 *A. quanzensis* individuals from 24 East African locations (APPENDIX 1 & 2). Fresh cambium or leaves were silica-dried in the field to avoid DNA fragmentation. Total DNA was extracted using the NucleoSpin plant kit (Macherey-Nagel, Düren, Germany) or the DNeasy 96 Plant Kit (QIAGEN, GMbH, Germany) for the fresh material. For herbarium material, a CTAB protocol was used (Doyle and Doyle, 1987).

### 2.3. Genotyping of SSRs and SNPs

Six microsatellite markers isolated from *A. bipindensis* were amplified in two PCR multiplexes in all samples according to a previously published protocol (Donkpegan et al. 2015). Amplified fragments were separated on an ABI 3730 sequencer (Applied Biosystems, Lennik, The Netherlands) and sized using the Genemapper software in comparison with the SYBR Safe (Invitrogen, Merelbeke, Belgium) size standard.

Genotyping by Sequencing (GBS) was performed for a subset of individuals (39 *A. africana* and 14 *A. quanzensis* individuals) at the Institute for Genomic Diversity and Computational Biology Service Unit at Cornell University (Ithaca, NY) according to a published protocol (Elshire et al. 2011). To select the best enzyme for the GBS protocol, one microgram of DNA of *Afzelia bipindensis* was used to build test libraries using three different enzymes: ApeKI (4.5-base cutter), EcoT22I and PstI (both 6-base cutters). Libraries were checked for appropriate fragment sizes (<500bp) and distribution on an Experion automatic electrophoresis system (Bio-Rad, Laboratories, CA, USA). The enzyme EcoT22I, giving appropriate fragment sizes (<500bp) was selected. To limit the risk of uneven coverage across loci and samples when applying GBS data to organisms with large genome sizes, we built and sequenced two independent libraries per individual, or pooled several DNA extracts per individual. Before library construction, DNA extracts were purified using a ZR-96 DNA Clean up kit (Zymo Research, Orange, CA), DNA quality was checked on a 1.5% agarose gel and DNA quantity was measured with Qbit HS (Invitrogen, USA). Overall, 95 GBS libraries were built corresponding to 154 DNA extractions obtained from 53 individuals of *Afzelia*. All libraries were sequenced together on one Illumina lane (HiSeq2000 San Diego, CA, USA), using 100-bp Single Read chemistry.

We used Sabre (https://github.com/najoshi/sabre) to demultiplex barcoded reads. After demultiplexing, sequence quality was evaluated with FastQC version 0.11.15 (Andrews, 2010). Low quality bases and adapter contamination were removed with TRIMMOMATIC version 0.33 (Bolger et al., 2014) with the following options: ILLUMINACLIP 2:30:10, LEADING 3, TRAILING 3, SLIDINGWINDOW 4:15, MINLEN 36.

First, a *de novo* assembly of GBS reads was carried out (including sequence reads of tetraploid African *Afzelia* species *A. bella*, *A. pachyloba* and *A. bipindensis*) using pyRAD v.3.0.2 software (Eaton & Ree, 2013) to produce a catalogue of GBS loci (3749 contigs, approximate length of 100bp per contig). This catalogue was used as a reference for mapping the reads of all individuals using BWA 0.7.5a-r405 (Li & Durbin 2009). The resulting alignments were converted to BAM format and reads were realigned around indels using SAMtools 0.1.17 (Li et al. 2009). The resulting BAM files were used as input for HaplotypeCaller algorithm of Genome Analysis Toolkit (GATK) v3.7 with standard parameters, to detect polymorphisms in each sample into a VCF format including SNPs and INDELs (DePristo et al. 2011). VCFtools (http://vcftools.sourceforge.net/) was used to remove indel variation, and retain only biallelic variants (SNPs) with ≤ 60% missing data within each species.

### 2.4. Data analysis

#### Population genetics parameters at geographic populations level

In order to characterize the diversity within each species at SSRs, we computed the allelic richness (*N*_*a*_), the effective number of alleles (*N*_*ae*_) following Nielsen et al. (2003), the observed heterozygosity (*H*_O_), the expected heterozygosity (*H*_E_), the inbreeding coefficient (*F*) and the genetic differentiation based on allele identity with the statistic *F*_ST_ using SPAGeDi 1.5a (Hardy, 2015). Permutation tests were used to test whether *F* or *F*_ST_ deviated from expectations of panmixia in SPAGeDi 1.5a (Hardy, 2015). For these analyses, we considered for both species, only populations sampled for a minimum of 5 individuals (Table 1). Null allele frequencies were estimated with INEST 1.0 (Chybicki & Burczyk, 2009), which also provided a corrected estimation of the inbreeding coefficient *F*. The selfing rate (*S*) was estimated in local populations with the largest sample sizes (samples ≥ 25 individuals – Table 1), based on the standardized identity disequilibrium assuming a mixed mating model (i.e. a proportion *s* of selfing and 1−*s* of random outcrossing) with standard errors (SE) estimated by jackknifing over loci (Hardy, 2015; David et al., 2007).

**Table 1.** Genetic diversity parameters and selfing rate estimates in populations of two *Afzelia* species. Number of genotyped trees (*N*), number of alleles per locus (N_a_), effective number of alleles (*N*_*ae*_), expected (*H*_E_) and observed (*H*_O_) heterozygosity, inbreeding coefficient estimated from heterozygote deficit (*F* = 1−H_O_/H_E_), inbreeding coefficient estimated while accounting for null alleles following the method implemented in INEst (*F*_*(null)*_). *p < 0.05 indicates significant deviation from HWE. NC indicates that no estimation was computed by INEst.

|  | Country | Populations | Latitude | Longitude | *N* | *N<sub>a</sub>* | *N<sub>ae</sub>* | *H<sub>E</sub>* | *H<sub>O</sub>* | *F* | *F<sub>(null)</sub>* | Selfing (*S*) |
| --- | --- | --- | --- | --- | --- | --- | --- | --- | --- | --- | --- | --- |
| *A. africana* | Benin | BassilaS | 2.4227 | 9.0037 | 7 | 3.00 | 2.53 | 0.459 | 0.595 | -0.327* | 0 | ∅ |
|  | Benin | BassilaN1 | 2.2886 | 9.0122 | 9 | 4.83 | 4.71 | 0.588 | 0.556 | 0.059 | 0 | ∅ |
|  | Benin | BassilaN2 | 2.2737 | 8.8221 | 11 | 4.33 | 4.28 | 0.516 | 0.470 | 0.094 | 0 | ∅ |
|  | Benin | BassilaN3 | 1.5556 | 9.2671 | 5 | 3.67 | 3.81 | 0.594 | 0.558 | 0.067 | 0 | ∅ |
|  | Benin | Lama | 2.1141 | 6.97694 | 34 | 5.00 | 2.45 | 0.497 | 0.434 | 0.129 | 0 | 0.33±0.16 |
|  | Benin | Natitingou | 1.3808 | 10.2785 | 9 | 4.50 | 3.78 | 0.561 | 0.444 | 0.218* | 0 | ∅ |
|  | Benin | ParcW1 | 11.2962 | 5.5864 | 18 | 5.17 | 3.69 | 0.589 | 0.574 | 0.026 | 0 | ∅ |
|  | Benin | ParcW2 | 1.2886 | 6.9592 | 6 | 3.17 | 3.17 | 0.483 | 0.528 | -0.105 | 0 | ∅ |
|  | Benin | Pendjari | 2.9909 | 11.5101 | 25 | 7.17 | 4.35 | 0.641 | 0.693 | -0.083 | 0 | 0 |
|  | Benin | Penessoulou1 | 3.0543 | 11.4788 | 32 | 5.67 | 3.30 | 0.483 | 0.458 | 0.052 | 0 | 0±0.1 |
|  | Benin | Penessoulou2 | 1.5242 | 10.9432 | 8 | 3.00 | 2.65 | 0.442 | 0.479 | -0.092 | 0 | ∅ |
|  | Togo | Notse | 1.5071 | 9.2856 | 12 | 4.17 | 2.99 | 0.525 | 0.528 | -0.005 | 0 | ∅ |
|  | Cameroon | Ngambetica | 1.6583 | 8.9988 | 7 | 3.83 | 3.08 | 0.566 | 0.500 | 0.125 | 0 | ∅ |
|  | Cameroon | Yoko | 12.0073 | 5.4309 | 7 | 3.33 | 2.47 | 0.500 | 0.524 | -0.052 | 0 | ∅ |
| *A. quanzensis* | Kenya | Gede | -3.3014 | 39.9965 | 31 | 5.50 | 3.45 | 0.473 | 0.409 | 0.139* | 0 | 0±0.07 |
|  | Kenya | Witu | -2.3837 | 40.5212 | 48 | 6.00 | 3.23 | 0.521 | 0.411 | 0.212* | 0 | 0±0 |
|  | DRC | Lubembe | -10.9166 | 22.5346 | 9 | 1.83 | 1.68 | 0.457 | 0.278 | 0.643* | NC | ∅ |

In order to characterize genomic diversity for each species for the GBS data, we computed nucleotide diversity Pi (π), corresponding to the average number of nucleotide differences per SNP site between pairs of sequences (Nei 1987), using DnaSP v. 5.10.01 software (Librado & Rozas 2009).

#### Population genetic structure

For SSR data, we used the Bayesian clustering method implemented in STRUCTURE 2.3.1 (Falush et al., 2003) to detect any putative genetic discontinuities within *A. africana* and *A. quanzensis* separately. We ran STRUCTURE 10 times for each number *K* of genetic clusters, using K=1–5. We ran 1,000,000 iterations after a burn-in period of 100,000 iterations, using the admixture model with independent allele frequencies between clusters, without considering the population of origin of each individual. We estimated LnP(K) and ΔK using the Evanno method (Evanno et al., 2005) implemented in STRUCTURE HARVESTER (Earl and vonHoldt 2012) to obtain the most likely value of K. We also used an alternative genetic clustering method implemented in the R package *tess3r* (Caye et al., 2016), which takes into account spatial information (the sampling location of each individual) to derive individual ancestry estimates. The default values of the program were used and each run (K◻=◻1–5) was replicated 10 times. The optimal value of K was defined by the minimum of the cross-entropy criterion.

For GBS-derived SNP data, we performed genetic clustering analysis using the sparse non-negative matrix factorization (sNMF) software, implemented in the R package LEA (Frichot et al., 2014). We also computed a genetic covariance matrix for each species to perform principal components analysis (PCA) using SMARTPCA (Patterson et al., 2006; Price et al., 2006) implemented in the SNPRelate package (Zheng et al., 2012).

#### Isolation by distance

Under Wright’s isolation-by-distance (IBD) model, the relatedness between individuals and/or populations is expected to decay linearly with the logarithm of their geographic distance on a two-dimensional scale (Hardy & Vekemans, 1999). To detect IBD within each species at large scales for SSR and SNP data, we calculated the kinship coefficient *F*_*ij*_ between individuals *i* and *j* using the estimator of Loiselle et al. (1995) implemented in SPAGeDi (Hardy and Vekemans, 2002). Positive and negative *F*_*ij*_ values indicate whether individuals are more related or less related than the average of two sampled individuals. Pairwise *F*_*ij*_ values were regressed on the logarithm of pairwise geographic distance, Ln(*d*_*ij*_), and IBD was tested by comparing the regression slope *b*_*log*_ to its distribution obtained from 10,000 permutations of the spatial locations of individuals. To illustrate IBD patterns, *F*_*ij*_ values were averaged over a set of distance classes (d) according to a geometric progression of eleven boundaries (0–1, 1–2, 2–5, 5–10, 10–50, 50–100, 100–200, 200–500, 500–1000, 1000–2000 and >2000 km) for *A. africana* and five (2–5, 5–10, 10–300, 300–500, >500 km) for *A. quanzensis* giving *F*(*d*). We used the *Sp*-statistic (Vekemans & Hardy, 2004) to quantify the strength of the spatial genetic structure: *Sp* = −*b*_*log*_/(1 − *F*_1_), where *F*_1_ is the mean *F*_*ij*_ between neighbouring individuals [approximated by *F*(d < 1 km) for the first distance class].

#### Demographic inference

Using SSR data, the demographic history of each species was assessed with the ‘bottleneck’ statistic T2 implemented in BOTTLENECK 1.2.02 (Piry et al., 1999). This statistic represents an average across loci of the deviation of the actual gene diversity *He* from the gene diversity expected from the number of alleles in the population assuming mutation-drift equilibrium in a population of constant size. If T2>0, the gene diversity excess reflects a loss of rare alleles possibly caused by recent founder events (bottlenecks), whereas population expansions almost always cause heterozygosity deficiency (T2<0, Cornuet & Luikart 1996). Simulations of the coalescent process were performed under three different mutation models: the infinite allele model (IAM), the stepwise mutation model (SMM), and the two-phase model with 70% of single-step mutations (TPM). The two latter models are considered to be more appropriate for SSR data. One thousand simulations were performed. Significant deviation from equilibrium gene diversity was determined using the Wilcoxon signed rank test, which is the most appropriate test when only few polymorphic loci are analysed (Piry et al., 1999).

For the SNPs data, to test for departure from the standard neutral model (SNM) the mean value of Tajima’s D (Tajima1989) over loci was computed and compared with the distribution of mean values from coalescent simulations using DnaSP v.5.10.1 (Librado & Rozas 2009). Tajima’s D statistic measures the standardized difference between nucleotide diversity π and the Watterson estimator θ per site (Watterson 1975). D is expected to be close to zero under the standard neutral model of population evolution, e.g., under a constant size population. High values of Tajima’s D suggest an excess of common variants, which can be consistent with balancing selection at the locus level, or with population contraction when detected at the genome level. Negative values of Tajima’s D, on the other hand, indicate an excess of rare variation, consistent with population growth when detected at the genome level, or with positive selection at the locus level (Tajima1989).

## 3. RESULTS

### SSR-based genetic diversity within each species and selfing rate

A total of 67 alleles were detected over all six loci for *A. africana* and the mean number of alleles per locus was 11.17 and ranged from 4 to 26 alleles. Observed and expected heterozygosity estimates per population ranged from *H*_*O*_ = 0.43 to 0.69 and from *H*_*E*_ = 0.44 to 0.64, respectively (Table 1). For *A. quanzensis*, a total of 42 alleles were detected over all six loci and the mean number of alleles per locus was 7.0 and ranged from 2 to 23 alleles. Observed and expected heterozygosity ranged from *H*_*O*_ = 0.28 to 0.41 and from *H*_*E*_ = 0.46 to 0.52. Inbreeding coefficients were not significantly different from zero in all populations (F = 0) after correcting for null alleles using INEST (APPENDIX 4). The estimated selfing rates *S* for three populations of *A. africana* (Lama, Penessoulou1 and Pendjari) and two of *A. quanzensis* (Gede and Witu) were close to zero (Table 1), except for Lama population (33%). F_*ST*_ statistics revealed low but statistically significant differentiation among populations, with weaker genetic structure in *A. africana*, *F*_*ST*_ = 0.045 (P < 0.01) than in *A. quanzensis*, *F*_*ST*_ = 0.078 (P < 0.01).

### GBS-based SNP data

After filtering to retain only biallellic SNPs, we obtained VCF files with 8541 SNPs for *A. africana* and 8730 SNPs for *A. quanzensis* using the GBS catalogue produced for the genus *Afzelia*. These files were then filtered to retain polymorphic SNPs within each species to remove SNPs and individuals with ≥ 60% missing data. After applying all filters, we removed nine individuals in *A. africana* and two in *A. quanzensis* and obtained VCF files containing 2800 polymorphic SNPs and 30 individuals in *A. africana* and 3841 polymorphic SNPs and 12 individuals in *A. quanzensis*. The final set of *A. africana* genotypes had an average missing data rate of 13.64% per sample with a mean depth of 40X. For *A. quanzensis*, the missing data rate was 28.48% per sample and 34X for mean depth. Total nucleotide diversity was π=0.00420 and θ=0.01124 in *A. africana*; π=0.03326 and θ=0.05094 in *A. quanzensis*.

### Population genetic structure

The STRUCTURE analyses of SSR data failed to detect population genetic structure at the intraspecific level. For both species, K◻=◻1 received the strongest support (APPENDIX 3). Runs assuming K=◻2 to K◻=◻5 revealed admixed ancestry of individuals with similar contributions of genetic clusters. The inclusion of geographic prior information using *tess3r* showed similar results, although *A. quanzensis* displayed somewhat uneven contributions of genetic clusters suggesting weak population substructure (APPENDIX 6). Conversely, the two SNPs data showed some evidence of genetic structure. The number of genetic clusters that best described the data was K=3 in *A. africana*, based on the criterion of minimum cross entropy (Figure 1, APPENDIX 7a). In *A. africana* only two gene pools occurred widespread across West Africa, without clear geographic pattern and many admixed individuals between these gene pools. The third gene pool was centred on Nigeria. The PCA shows low levels of genetic differentiation (variance explained by PC1 and PC2 are 7.10% and 5.90%, respectively) and highlights the divergence of the Nigeria cluster (Figure 1C). In *A. quanzensis*, cross entropy values decreased with increasing K up to the maximum number of K tested (K=10) suggesting a stronger population genetic structure in this species (APPENDIX 7b). This is confirmed by the PCA (PC1 and PC2 explain 18.32% and 12.79% of the variance, respectively, Figure 1D). We chose to retain K=2 to represent the highest hierarchical level of genetic structure; higher values of *K* revealed evidence for additional genetic structuring in the sample. One cluster covered the north-eastern part of the sample range and was mostly represented in coastal Kenya whereas the other one was widespread across the sample range (Fig 1).

**Figure 1.**
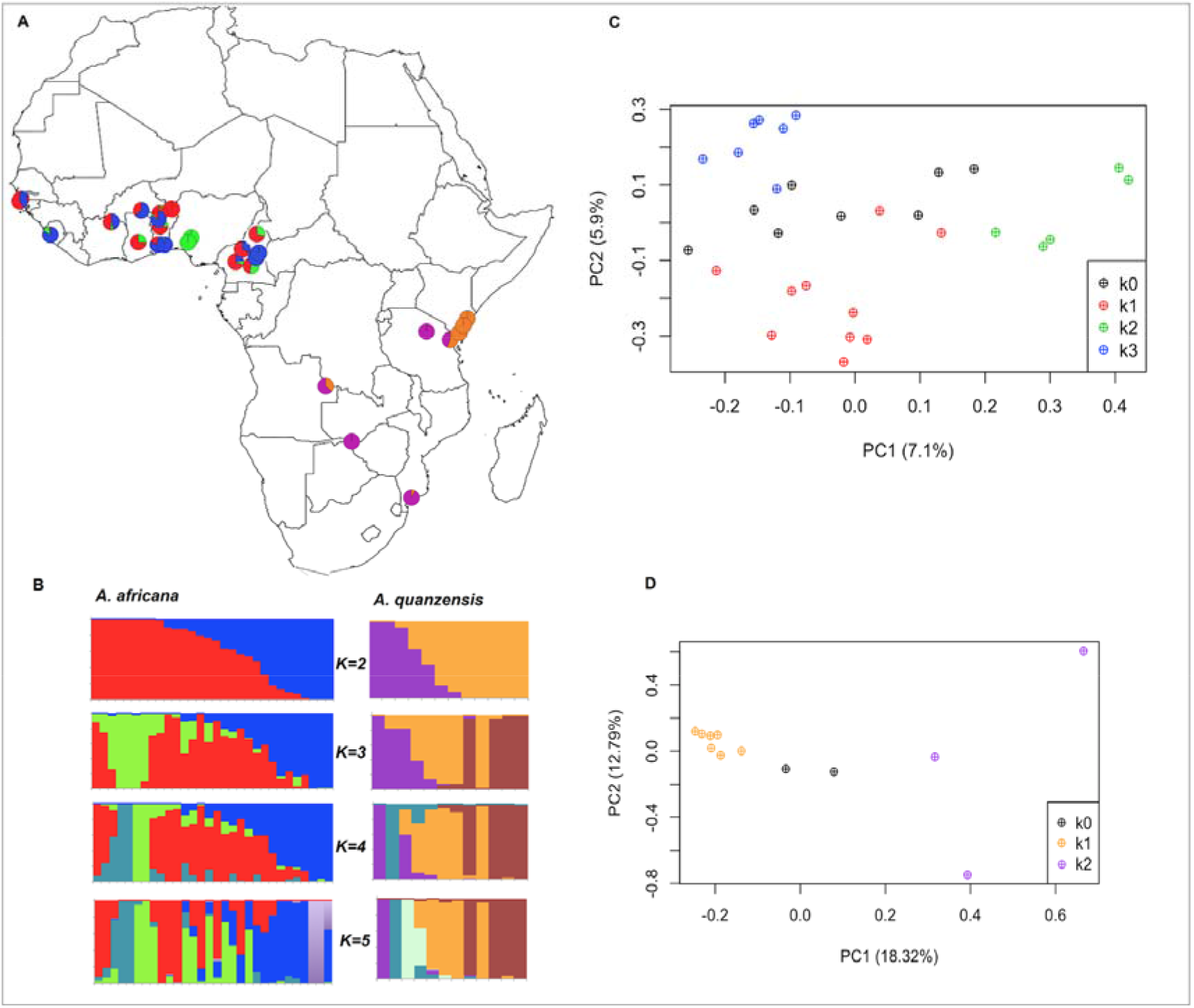
Genetic structure of African diploid *Afzelia* species using GBS-based SNPs (N = 30 *A. africana* with 2800 SNPs; N = 12 *A. quanzensis* with 3841 SNPs). **A)** Geographic origin of samples and population genetic structure of *A. africana* at K=3 (Western Africa) and of *A. quanzensis* at K=2 (East and Austral Africa), where pie charts represent individual ancestry proportions in the assumed populations, as estimated using sNMF. **B)** Histograms of individual ancestry proportions for each species, as estimated using sNMF for K=2 to K=5 assumed ancestral populations. **C - D)** PCA ordinations along the first two PCA axes of (**C**) *A. africana* and (**D**) *A. quanzensis*, where symbols distinguish sNMF clusters (k0 represent samples not assigned to a cluster at q>0.7).

### Patterns of isolation by distance (IBD)

Pairwise kinship declined with increasing geographic distance for both types of markers (Figure 2). In both species, kinship for the first distance class (ca. 1000 m for *A. africana* and 5000 m for *A. quanzensis*) ranged around 0.05 for SSRs and 0.06 for SNPs, and quickly dropped with distance, indicating a signature of isolation by distance. The regression slope *b*_*Ld*_ was significantly negative in *A. africana* (*b*_*Ld*_= −0.005 for SSRs and −0.008 for SNPs; both *P* < *0.001*) and *A. quanzensis* (*b*_*Ld*_= −*0.007* for SSRs and −0.006 for SNPs respectively; both *P* < *0.001*). Overall, similar patterns of IBD were detected for both types of markers in both species.

**Figure 2.**
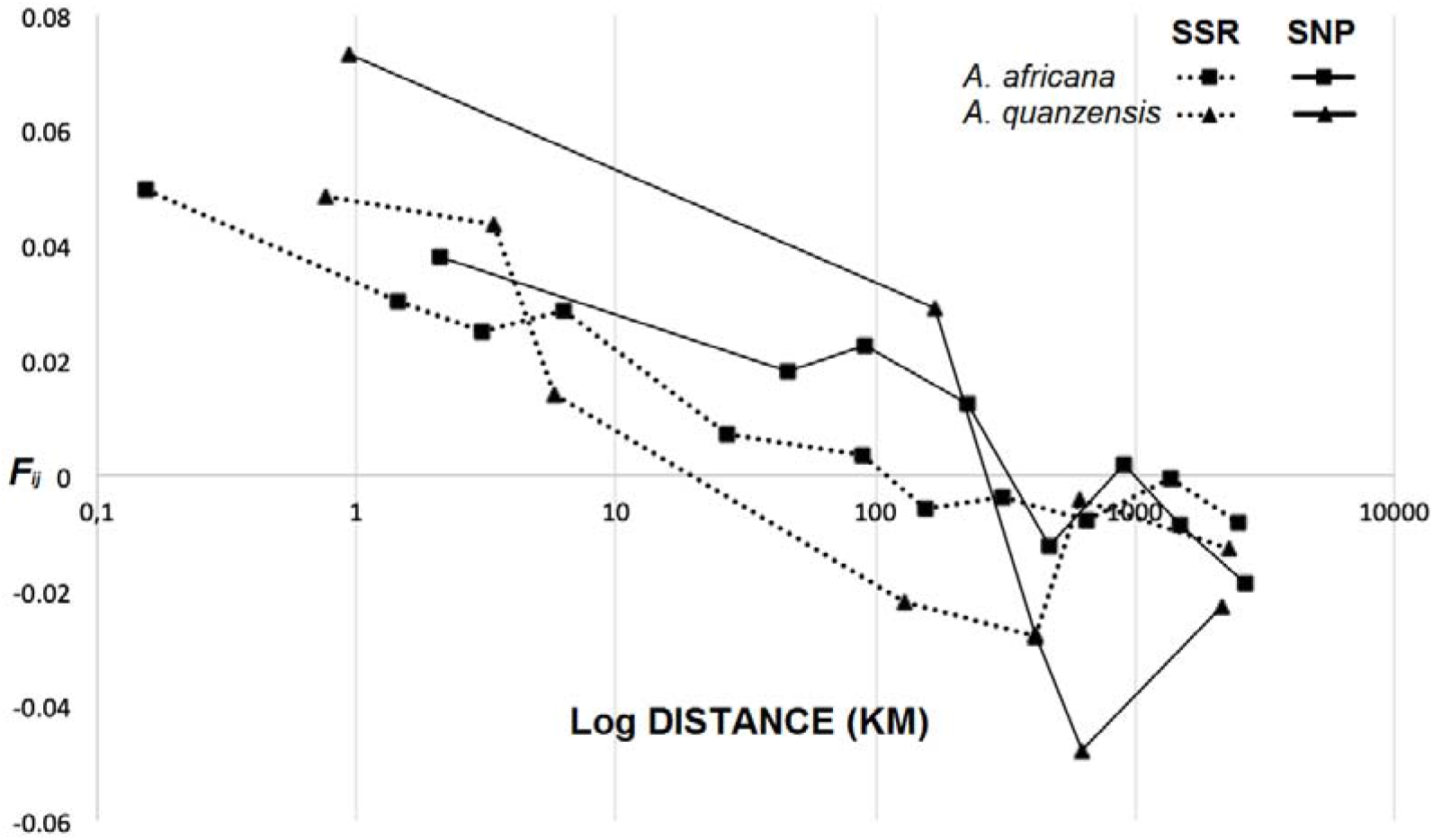
Spatial genetic structures (kinship-distance curves) of *A. africana* (square) and *A. quanzensis* (triangle) based on SSRs (stippled lines) and SNPs (plain lines).

### Demographic inference in each species

With SSRs, under the three models implemented in BOTTLENECK (IAM, TPM and SMM), both species showed a negative value of T2 and a significant heterozygosity deficiency (P<0.01) for *A. africana* and for *A. quanzensis* after Wilcoxon tests (Table 2). These results suggest absence of a recent bottleneck at the species level for both species.

**Table 2.**
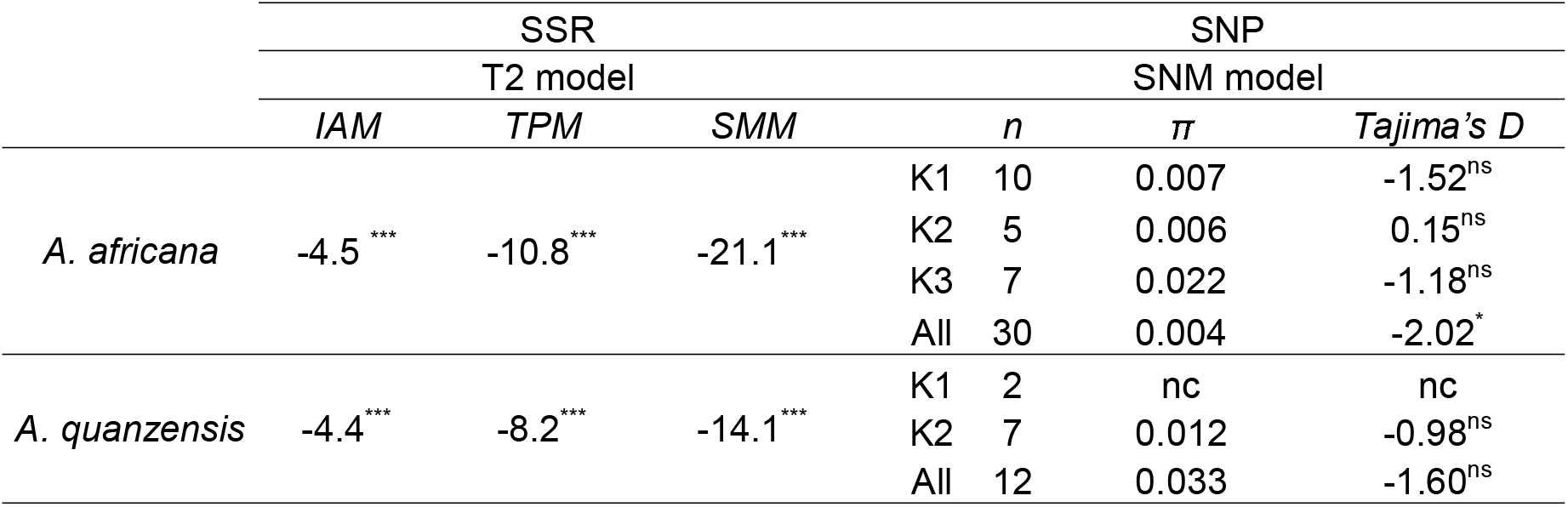
An evaluation of alternative demographic models for the total population of both species with SSR and SNP. T2 is the bottleneck statistic of different models; IAM, infinite allele model; TPM, two-phase model; SMM, stepwise mutation model; SNM, standard neutral model; n, number of individuals; π, nucleotide diversity and K1, K2 and K3 represent the genetic groups defined for each species (see Figure 1). nc, not computed; ns: not significative; * P<0.05; ** P<0.01; *** P<0.001

For GBS data, mean Tajima’s D estimates were negative in both species, with values of −2.017 (P<0.05) for *A. africana* and −1.598 (not significant) for *A. quanzensis*. These results are in agreement with a signature of population expansion in *A. africana* at the species level, whereas in *A. quanzensis*, the standard neutral model of constant species-level population size could not be rejected (Table 2).

## 4. DISCUSSION

### Large-scale population structure

Our results reveal a pattern of isolation by distance in the two savannah representatives of the genus *Afzelia* in Africa -*A. africana* and *A. quanzensis*-, i.e., the kinship between individuals decreased with spatial distance. It is worth noting that SSRs and SNPs gave very similar IBD patterns despite large differences in the number of loci and sampling strategies, as observed in previous studies (Yang et al., 2011). The IBD observed probably reflects the limited movement of pollen and seeds. However, the mechanisms of pollination and seed dispersal are not well known in *Afzelia*. The local movement of seeds would be expected given the fact that the seeds of *Afzelia* are heavy and also given the observation that small rodents act as dispersers (*Cricetomys emini, Epixerus wilsoni, Protoxerus stangeri*, Bationo et al., 2001; Evrard, 2015). However, long-distance seed dispersers such as monkeys (*Cercopithecus albogularis*) and birds -mainly hornbills- (Van Wyk & Van Wyk, 1997; Gathua, 2000) have been also reported. The pollination mechanism is even less studied. While large *Xylocopa* bees (Kato et al., 2008) act as pollination agents in Asian *Afzelia*, their African congeners *Apis mellifera scutellata* would not be able to transfer pollen beyond 3.2 km (Dick et al., 2003).

While SSRs could not retrieve distinct genetic clusters across the natural range of *A. africana* in the Sudanian savannah and *A. quanzensis* in the Zambezian savannah, SNP data revealed genetic groups within species (particularly more pronounced *in the latter*). However, the genetic clusters identified by SNPs exhibit high levels of admixture and do not correspond to any clearly delimited geographic entities. This structure might thus reflect solely the trend of IBD rather than a history of past population fragmentation. These observations suggest that gene flow has been restricted but populations have remained connected throughout the large, continuous Sudanian or Zambezian savannahs. The higher discriminating power of SNPs over SSRs for detecting genetic clusters has also been reported previously (e.g. Liu et al 2005; Fischer et al., 2017).

Different scenarios were tested to reconstruct the demographic history of each species. SSR and SNP data were again congruent in detecting signatures of population expansion. However, our data were not powerful enough to identify if these signatures reflect range expansions (and from which source) or only a demographic expansion without change of distribution. In any case, populations of both savannah species apparently did not experience major disturbances leading to their fragmentation as has been suggested in some other savannah species (Bryja et al. 2010; Odee et al. 2012, Sexton et al., 2015).

### Comparison with other tropical trees in Africa

our results reveal no cut clear genetic discontinuities over large distances in the Sudanian and the Zambezian savannahs for *A. africana* and *A. quanzensis*, respectively, and are consistent with those observed in other savannah tree species, namely *Adansonia digitata* and *Khaya senegalensis*, which showed no geographic discontinuities of the genetic variation, and the moderate levels of differentiation found in *Acacia senegal*. These results suggest that the African savannahs have not experienced major upheavals leading to their fragmentation (Salzmann et al., 2002; Vincens et al., 2006; Watrin et al., 2009) in contrast to the major fluctuations of the rainforest cover over time (Maley, 1996). The cases of *Vitellaria paradoxa* (Allal et al., 2011; Logossa et al., 2011) and *Parkia biglobosa* (Lompo et al., 2018), which show different geographic and genetic clusters in the Sudanian region (but include large genetically homogenous clusters in central west Africa) might be due to their high socio-economic importance in agroforestry systems in savannah parklands because they all produce seeds that are marketed and widely used in human food. Whether their genetic structures have been influenced by human activities remains an open question.

In the last few years population genetic data have accumulated for a number of African rainforest trees, indicating strong differentiation of the tree populations in Central and West African rainforests for most of the tree species (Hardy, Born et al. 2013, Heuertz, Duminil et al. 2014).This genetic structuring cannot be explained by current geographic barriers such as the main mountain chains (Cameroonian Volcanic Line, Cristal Mountains, and Chaillu massif) or major rivers in the region (Sanaga, Dja, and Oougué river). Molecular dating suggests historical isolation of the tree populations, probably led by rainforest fragmentation, during the cold and dry Ice-Age periods of the Pleistocene (<2.58 Myra). These results contrast with the genetic connectivity found for the *Afzelia* and other savannah tree species over large Sudanian and Zambezian ranges.

### Local-scale genetic diversity with SSRs

Inbreeding and selfing rates remain very low in adult populations of *A. africana* and *A. quanzensis*. Gene diversity parameters for SSRs markers showed a large range of local genetic diversity in our study (*A. africana* : *H*_*E*_ = 0.46–0.66 and *A. quanzensis*: *H*_*E*_ = 0.40–0.66) and in other population-level studies of *A. quanzensis* from Zimbabwe (*H*_*E*_ = 0.41–0.51; Jinga & Ashley, 2018), *A. africana* from Benin (*H*_*E*_ = 0.09–0.88; Houehanou et al., 2019), and the Asian congener *A. xylocarpa* (*H*_*E*_ = 0.47–0.66; Packkad et al., 2014). Comparable genetic diversity ranges were documented in the investigated African savannah tree species *Khaya senegalensis* (*H*_*E*_ = 0.44–0.71; Sexton et al., 2015), *Vitellaria paradoxa* (*H*_*E*_ = 0.42–0.62; Allal et al., 2011), *Acacia senegal* (*H*_*E*_ = 0.63–0.70; Omondi, et al., 2010) and *Parkia biglobosa* (*H*_*E*_ = 0.61–0.82; Lompo et al., 2018). Much lower levels were documented in *Adansonia digitata* (*H*_*E*_ = 0.27–0.35; Kyndt et al., 2009). Despite the high influence of past climate changes and signature of forest fragmentation on rainforest tree species no remarkably lower population genetic diversity is observed: *Aucoumea klaineana* (*H*_*E*_ = 0.38–0.55; Born et al., 2008), *Milicia excelsa* (*H*_*E*_ = 0.53–0.56; Bizoux et al., 2009), *Baillonella toxisperma* (*H*_*E*_ = 0.56–0.58; Ndiade-Bourobou et al., 2010), *Distemonanthus benthamianus* (*H*_*E*_ = 0.47–0.58; Debout et al., 2011), *Greenwayodendron suaveolens* (*H*_*E*_ =0.7-0.8; Piñeiro et al., 2017), *Scorodophloeus zenkeri* (*H*_*E*_ =0.50-0.60; Piñeiro et al., 2017), *Terminalia superba* (*H*_*E*_ = 0.51–0.81; Demenou et al., 2018).

## 5. CONCLUSION

The SSR and SNP-based data analyses of *Afzelia* species from the African savannahs have shown that both species did not exhibit strong geographic barriers to genetic connectivity across their Sudanian and Zambezian ranges, although isolation by distance patterns indicate restricted gene flow. In this study, both markers provided overall congruent results, although SNP had more resolution power than SSRs for population genetic structure analyses. Demographic analyses with both SNPs and SSRs data suggested demographic expansion. Collectively, these data demonstrate the strong influence that savannah ranges exert on genomic diversity, within across their population range. Thus, there is consistent evidence for the signature of population expansion beginning to accumulate in the genome of these savannah species; in contrast to forest species, which show a long history of fragmentation in most of studied species in Guineo-Congolian rainforest (Hardy et al., 2013).

## ACKNOWLEDGMENTS

This work received financial support from the “Fonds pour la Formation à la Recherche dans l’Industrie et l’Agriculture (FRIA-FNRS, Belgium)” through a research grant to A.D., from the Marie Curie FP7-PEOPLE-2012-IEF program (project AGORA) awarded to R.P., from the Fonds de la Recherche Scientifique (F.R.S.-FNRS) through project J.0292.17F, the Belgian Science Policy (project AFRIFORD) and the CGIAR Research Program on Forests, Trees and Agroforestry. This work has benefited from support of a grant from Investissement d’Avenir grants of the ANR (CEBA:ANR-10-LABX-25-01). A.D. acknowledges a Labex COTE Mobility grant to INRA. The authors are grateful to Nils Bourland who helped us during field expeditions, through the project PD 620/11 Rev.1 (M): “Development and implementation of species identification and timber tracking in Africa with DNA fingerprints and stable isotopes” by the International Tropical Timber Organization (ITTO). We also thank the Botanic Garden of Meise (BR-Herbarium, Belgium), ULB (BRLU-Herbarium) and Naturalis (WAG-Herbarium, Netherlands) who provided us with material from their herbarium collections; and Esra Kaymak and Tom Gilbert for their assistance in generating GBS data.

## DATA ACCESSIBILITY

Microsatellite and GBS data are being submitted in DRYAD and GenBank’s Sequence Read Archive respectively.

## AUTHOR CONTRIBUTIONS

A.D., J-L.D. and O.H. conceived the study. A.D. collected the data and performed the analyses. R.P. generated the GBS data sequencing. A.D., R.P., M.H., J.D., K.D., J-L.D., O.H. interpreted the results, contributed to drafting and writing the article.

