## Supporting information for "Population genomics of the widespread African savannah trees *Afzelia africana* and *Afzelia quanzensis* (Caesalpinioideae, Fabaceae) reveals no significant past fragmentation of their distribution ranges"

**APPENDIX 1: Table S1.** Population’s origins of fresh plant tissue samples used for the SSRs analyses of *Afzelia* savannah species (N = sample size per population)

**APPENDIX 2: Table S2.** Herbarium material used for the SSRs analyses of *Afzelia* savannah species

**APPENDIX 3: Table S3.** Mean Log-likelihood for each number of genetic clusters (K) obtained from the use of STRUCTURE software on the SSR data of *Afzelia*.

**APPENDIX 4: Table S4.** Estimated proportion of null alleles per SSR locus in *Afzelia* species according to INEst software.

**APPENDIX 5: Figure S1.** Distribution map of *A. africana* (triangles) and *A. quanzensis* (stars) samples analysed and their location in African savannah biogeographic zones delineated by Linder et al. (2012)

**APPENDIX 6: Figure S2.** Genetic structure of African diploid *Afzelia* species using SSRs (N = 241 *A. Africana*; N = 113 *A. quanzensis*), using the tess3r software. The ancestry proportions inferred in each of K ancestral populations (different colours) are shown for each individual, represented by a vertical bar.

**APPENDIX 7a: Figure S3a.** The number of K (1-10) ancestry components best explaining the genetic structure of *A. africana* assessed using the cross-entropy criterion obtained from the sNMF program on SNP data. The best-fitting model (i.e., the model with the lowest minimal cross-entropy) had three ancestry components (K = 3) identified for the 30 individuals used with 2800 polymorphic SNPs.

**APPENDIX 7b: Figure S3b.** The number of K (1-10) ancestry components best explaining the genetic structure of *A. quanzensis* assessed using the cross-entropy criterion obtained from the sNMF program on SNP data. The best-fitting model (i.e., the model with the lowest minimal cross-entropy) is not clearly identified for the 12 individuals used with 3841 polymorphic SNPs.

**APPENDIX 1: Table S1.**

| **Species** | **Country** | **Population** | **Latitude** | **Longitude** | **N** |
| --- | --- | --- | --- | --- | --- |
| *A. africana* | Benin | Bassila | 1.556 | 9.263 | 32 |
|  |  | Lama | 2.135 | 6.977 | 34 |
|  |  | ParcW | 2.992 | 11.503 | 24 |
|  |  | Pendjari | 1.525 | 10.932 | 25 |
|  |  | Pénéssoulou | 1.556 | 9.267 | 40 |
|  | Burkina-Faso | Comoé | 4.593 | 9.915 | 2 |
|  |  | Pama | 0.782 | 11.251 | 3 |
|  | Cameroon | Ngambetica | 11.296 | 4.789 | 7 |
|  |  | Sibati | 13.666 | 5.267 | 7 |
|  |  | Yoko | 12.314 | 5.431 | 2 |
|  | Senegal | Casamance | -16.002 | 12.799 | 4 |
|  | Togo | Notsé | 1.289 | 6.958 | 12 |
| *A. quanzensis* | Kenya | witu | -2.384 | 40.521 | 49 |
|  |  | Gede | -3.301 | 39.996 | 31 |
|  |  | kwalé | -4.183 | 39.440 | 3 |
|  | DRC | Lubembe | -10.917 | 22.535 | 10 |
|  | Zimbabwe | Victoria falls | -17.933 | 25.833 | 1 |

**APPENDIX 2: Table S2.**

| **Species** | **Geographic origin** | **Herbarium** | **Collectors** | **Voucher** |
| --- | --- | --- | --- | --- |
| *A. quanzensis* | Mozambique | WAG | na | WAG1704773 |
|  | Mozambique | WAG | na | WAG1704771 |
|  | Mozambique | WAG | na | WAG1704769 |
|  | Mozambique | WAG | na | WAG1704765 |
|  | Mozambique | WAG | na | WAG1704764 |
|  | Mozambique | WAG | na | WAG1704757 |
|  | Tanzanie | WAG | na | WAG1704751 |
|  | Tanzanie | WAG | na | WAG1704749 |
|  | Tanzanie | WAG | na | WAG1704744 |
|  | Afrique du sud | WAG | na | WAG1704737 |
|  | RDC | WAG | Duvigneaud & Timperman | 2496 |
|  | Angola | WAG | R. Dechamps & F. Kurta | 1302 |
|  | Mozambique | WAG | de Koning Jan | 8109 |
|  | Mozambique | WAG | Groenendi E.M.C. | 266 |
|  | Mozambique | WAG | Van de Koning | 9575 |
|  | Tanzanie | WAG | J.C. Lovett | 4915 |
|  | Tanzanie | WAG | Mwangoka, M.A. | 350 |
| *A. africana* | RCA | BRLU | Tisserant | 3046 |
|  | RCA | BRLU | Tisserant | 2407 |
|  | Cameroun | BRLU | de Wilde W.J.J.O. | 2793 |
|  | Ouganda | BRLU | R.G. Sangstes | S. 223 |
|  | Guinea-Bissau | BRLU | Verjans J.-M. | na |
|  | Guinea-Bissau | BRLU | Malaisse F. | na |
|  | Guinnea-Bissau | BRLU | Kasper & Descheres | na |
|  | Cameroun | BR | C. Geerling & J. Nene | 4708 |
|  | Cameroon | BR | Geerling, C. | 4708 |
|  | Cameroon | BR | Wit, P. | 2944 |
|  | Cameroon | BR | Zon, A.P.M. van der | 2579 |
|  | Cameroon | BR | Wilde, W.J.J.O. de | 2793 |
|  | Ghana | BR | Jongkind, C.C.H. | 2510 |
|  | Ghana | BR | Schmidt, H.H. | 1782 |
|  | Ghana | BR | Morton, J.K. | 1918 |
|  | Ghana | BR | Cudjoe, F.S. | 623 |
|  | Ghana | BR | Jongkind, C.C.H. | 1963 |
|  | Ghana | BR | Schmidt, H.H. | 3370 |
|  | Ivory Coast | BR | Geerling, C. | 2044 |
|  | Ivory Coast | BR | Versteegh, C. | 521 |
|  | Ivory Coast | BR | Wilde, W.J.J.O. de | 783 |
|  | Togo | BR | Thijssen, M.T. | 222 |
|  | Ivory Coast | BR | Wilde, J.J.F.E. de | 3474 |
|  | Ivory Coast | BR | Versteegh, C. | 325 |
|  | Ivory Coast | BR | Oldeman, R.A.A. | 332 |
|  | Ivory Coast | BR | Jongkind, C.C.H. | 4410 |
|  | Nigeria | BR | Chapman, J.D. | 2752 |
|  | Nigeria | BR | Geerling, C. | 3122 |
|  | Nigeria | BR | Leeuw, P.N. de | 70 |
|  | Burkina Faso | BR | Küppers, K. | 1036 |
|  | Sierra Leone | BR | Pollard, B.J. | 1405 |
|  | Siera Leone | BR | Cole, E.A. | 121 |
|  | Guinea-Bissau | BR | Espirito Santo, J.V.G. do | 1508 |
|  | Guinea-Bissau | BR | Espirito Santo, J.V.G. do | 3469 |
|  | Senegal | BR | Adam, J.-.G. | 8685 |
|  | Ghana | BR | Jongkind, C.C.H. | 2440 |

**APPENDIX 3: Table S3.**

|  | | *A. africana* | | *A. quanzensis* | |
| --- | --- | --- | --- | --- | --- |
| K | Reps | Mean LnP(K) | Stdev LnP(K) | Mean LnP(K) | Stdev LnP(K) |
| 1 | 10 | -4002.9 | 0.7 | -1607.3 | 1.8 |
| 2 | 10 | -4221.9 | 56.2 | -1740.9 | 54.5 |
| 3 | 10 | -5885.3 | 904.0 | -1975.4 | 304.8 |
| 4 | 10 | -4215.0 | 203.6 | -1892.1 | 419.5 |
| 5 | 10 | -7510.4 | 8117.3 | -2139.3 | 235.3 |
| 6 | 10 | -4558.1 | 192.1 | -2932.4 | 1572.9 |

**APPENDIX 4: Table S4.**

|  | *A. quanzensis* | | *A. africana* | |
| --- | --- | --- | --- | --- |
| Locus | *p_ij_* | SE | *p_ij_* | SE |
| R9-48_Q4 | 0.61 | 0.04 | 0.25 | 0.03 |
| R9-01_Q1 | 0.03 | 0.03 | 0.01 | 0.02 |
| R9-07_Q4 | 0.14 | 0.05 | 0.03 | 0.03 |
| R9-19_Q2 | 0.28 | 0.04 | 0.24 | 0.03 |
| R9-65_Q1 | 0.00 | 0.005 | 0.11 | 0.03 |
| R9-61_Q2 | 0.02 | 0.04 | 0.00 | 0.00 |

**APPENDIX 5: Figure S1.**

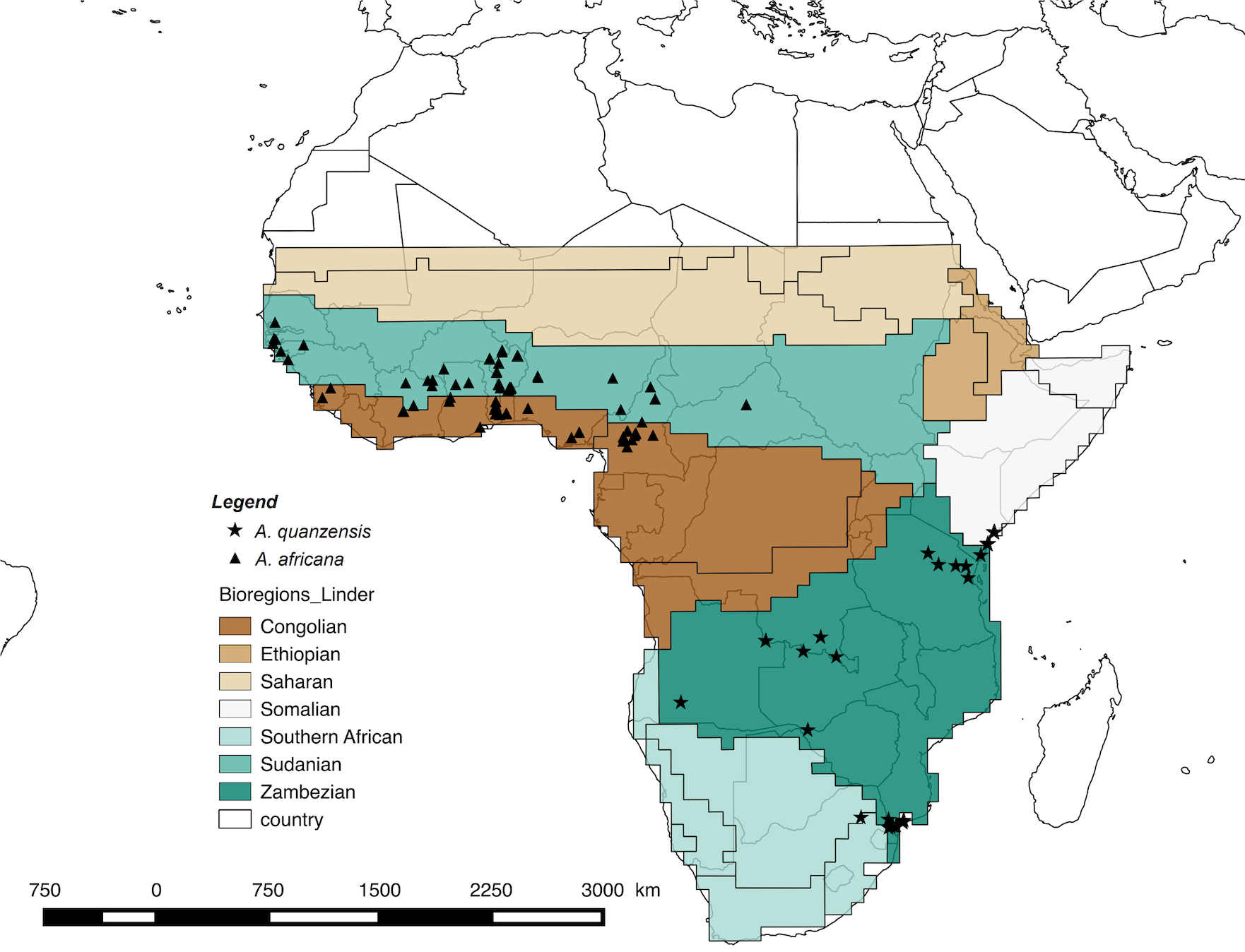

**APPENDIX 6: Figure S2.**

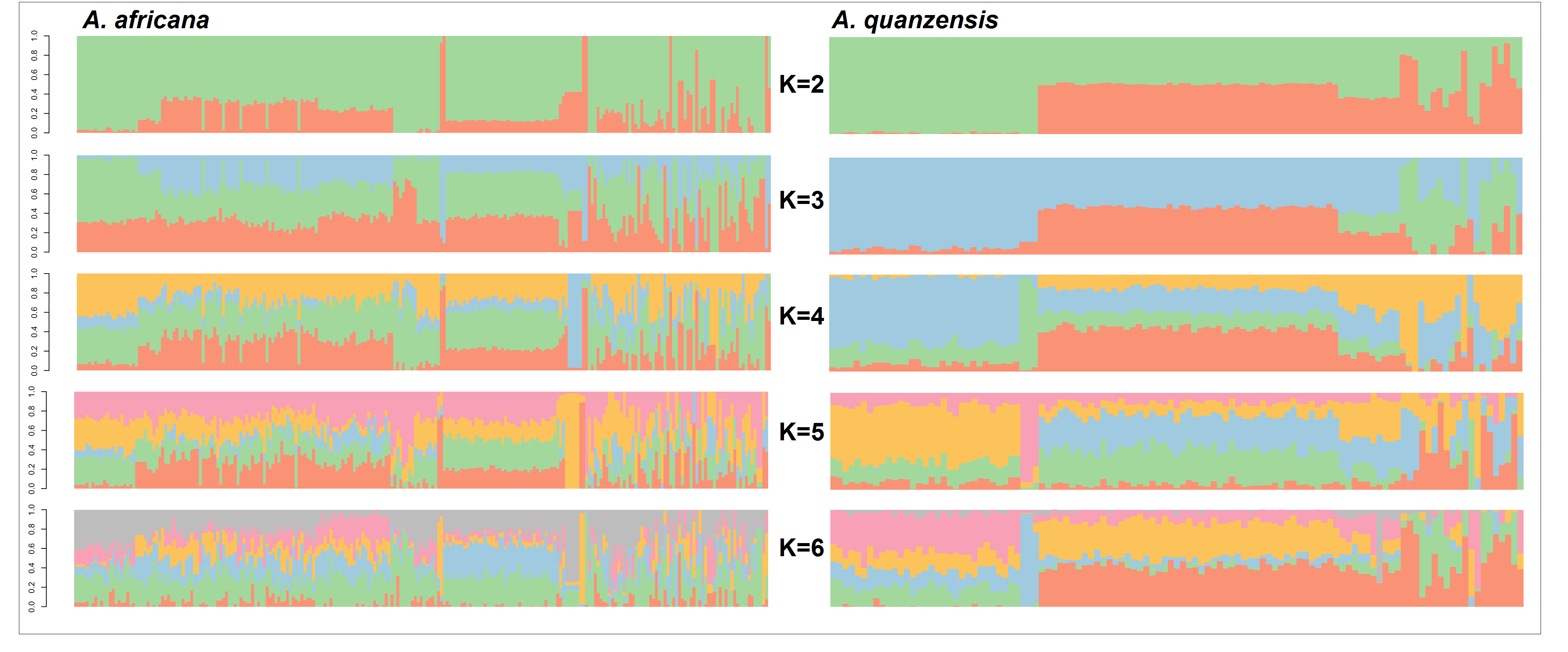

**APPENDIX 7a: Figure S3a.**

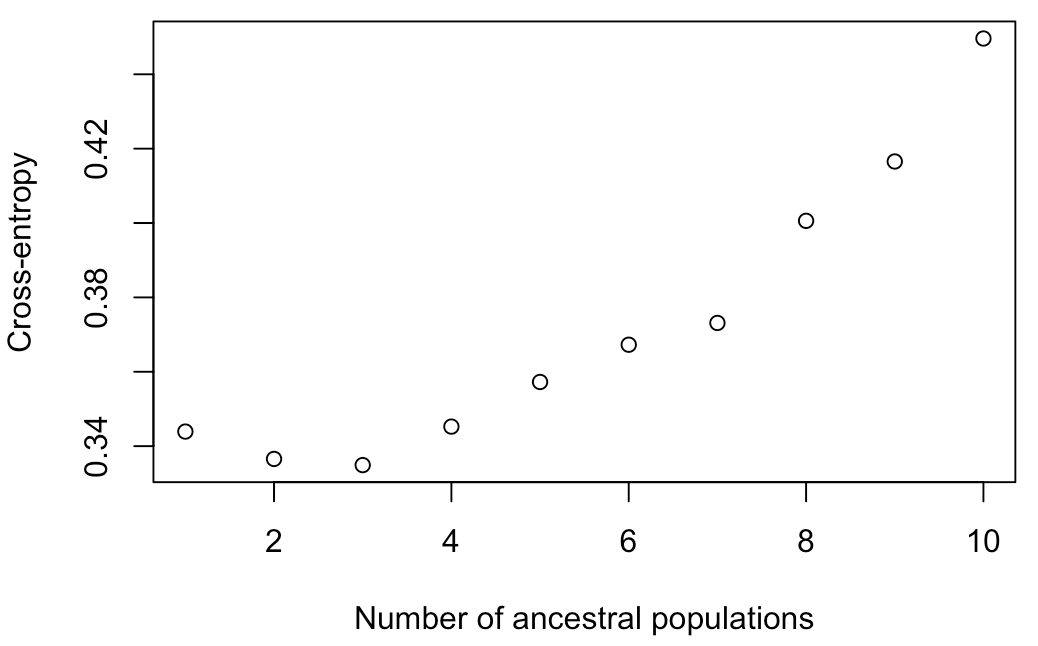

**APPENDIX 7b: Figure S3b.**

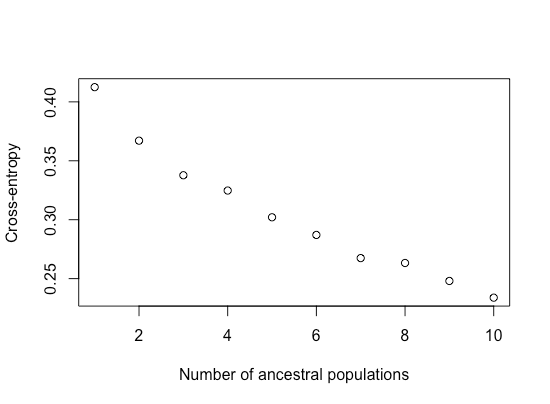
